# The effects of habits on motor skill learning

**DOI:** 10.1101/338749

**Authors:** Nicola J. Popp, Atsushi Yokoi, Paul L. Gribble, Jörn Diedrichsen

**Author notes:** Correspondence address Jörn Diedrichsen, The Brain and Mind Institute, University of Western Ontario, London, Canada.

## Abstract

Skill learning involves the formation of stable motor patterns. In musical and athletic training, however, these stable motor habits can also impede the attainment of higher levels of performance. We developed an experimental paradigm to induce a specific motor pattern in the context of a discrete sequence production task and to investigate how these habits affect performance over a 3-week training period. Participants initially practiced small segments of 2 to 3 finger movements (“chunks”) and then learned longer sequences composed of these chunks. This initial training induced a persistent temporal pattern during execution, with shorter inter-press-intervals within a chunk and longer ones at chunk boundaries. This pattern remained stable during the subsequent 10 days of training, in which participants were asked to produce the sequence as fast as possible from memory. The habit was also preserved when the sequences were directly displayed, removing the need for memory recall. We were able to induce chunking patterns that were either beneficial or detrimental to performance by taking into consideration the biomechanical constraints of the sequences. While we observed an overall reduction in the detrimental effect of the disadvantageous chunking instructions with training, our results show that the degree to which these detrimental chunk structures were maintained, was predictive of lower levels of final performance. In sum, we were able to induce beneficial and detrimental motor habits in a motor sequence production task and show that these initial instructions influenced performance outcomes over a prolonged period of time.

**Significance Statement:** A habit is defined as an automatized action that resists modification once sufficiently established. Preventing bad habits, while reinforcing good habits, is a key objective when teaching new motor skills. While habit formation is an integral part of motor skill acquisition, previous research has focused on habit formation in terms of action selection. In this paper, we examine habit formation in terms of motor skill execution, after the action has been selected. We were able to induce beneficial or detrimental motor habits in the production of motor sequences. Habits were stable over a prolonged training period. Our results demonstrate how cognitive instruction can lead to persistent motor habits and we explore how these habits are potentially modified with training.

## Introduction

What does it take to become an expert at a motor skill such as playing the piano? Clearly, practice is key. Some have proposed that 10,000 hours of training are necessary to develop a high level of performance (Ericsson et al., 1993; Hayes, 2013). However, simply practicing for many hours may not lead to expert performance, as numerous examples have shown (Haith and Krakauer, 2018). This is sometimes attributed to the formation of habits: automatic (Hélie, Waldschmidt, & Ashby, 2010; Moors & De Houwer, 2006) and highly entrenched behavioral patterns that resist change through retraining (Ashby et al., 2003; Jager, 2003; Seger and Spiering, 2011; Graybiel and Grafton, 2015; Hardwick et al., 2019).

Animal models have been integral to the study of habit formation and its neural underpinnings (Jog et al., 1999; Wickens et al., 2007; Smith and Graybiel, 2014, 2016; Robbins and Costa, 2017). However, the majority of animal experiments investigating habit formation have focused on habits in the context of action selection – i.e. choosing *what* action to perform. In contrast, in this paper we address the question of habits in motor performance – i.e. habits that influence *how* to perform a chosen action. For example, a tennis player could be influenced by a habitual pattern in action selection, whereby she always chooses a forehand over a backhand to return a serve. At the same time, she could be influenced by a motor habit, whereby she executes the forehand without rotating her hips.

Critical to the definition of a habit is that the behavior is maintained even though it is no longer adaptive (Adams, 1982; Dickinson, 1985; Dezfouli and Balleine, 2012).

Most experiments, therefore, demonstrate the existence of a habit by teaching subjects a behavior under one reward contingency and show its persistence when the reward contingency switches (Ashby et al., 2003; Smith and Graybiel, 2013a).

To investigate the influence of habit formation on motor skill learning we used a discrete sequence production task (DSP) in which participants performed an explicitly learned series of finger presses as fast as possible (Verwey, 2001; Abrahamse et al., 2013). Learning in this task depends on both cognitive and motor processes (Diedrichsen & Kornysheva, 2015; Wong, Lindquist, Haith, & Krakauer, 2015). Initial performance relies on forming a declarative memory of the sequence that can be sculpted through explicit instructions (de Kleine & Verwey, 2009; Verwey, Abrahamse, & Jiménez, 2009) and potentially can constrain subsequent motor optimization (Bo and Seidler, 2009; Seidler et al., 2012). We tested the hypothesis that the initial instruction causes the formation of a motor habit which influences the learning of execution-related skills in subsequent motor training.

We instructed participants to memorize long sequences of finger presses by first practicing a set of smaller 2-3 digit “chunks” on an isometric keyboard-like device (Miller, 1956; Verwey, 1996; Verwey and Dronkert, 1996; Halford et al., 1998; Wymbs et al., 2012). Two different chunk sets were used. Participants were then trained on seven 11-digit sequences. Each sequence was subdivided into chunks (depending on chunk set) so that boundaries between chunks were either aligned or misaligned with biomechanically easy or difficult finger transitions. This manipulation influenced initial performance with sequences learned using the aligned chunk structure being performed faster. After the introduction phase, participants had to recall the sequences from memory and practiced them over the course of 3 weeks.

We investigated three questions: First, do the initial instructions lead to a stable motor performance pattern and how long does it persist? Second, to what degree are these patterns maintained even if they are detrimental to performance? Finally, what learning-related changes are involved in overcoming motor habits?

## Methods

### Participants

Forty participants who reported no neurological conditions were recruited for the study (30 females; ages: 19 to 33). Thirty-two of them were randomly assigned to learn the sequences with one of the two chunk sets (Figure 1) and the remaining eight participants were assigned to a control group. All participants were right-handed based on the Edinburgh Handedness Inventory and completed informed consent. On average, participants had received 4.68 (± 5.55) years of musical training, with 55% percent reported having more than 6 months of experience playing the piano. While participants with piano experience performed the sequences faster than participants with no experience and the number of practice years correlated with execution speed (MT), the amount of participants’ prior musical experience did not have a qualitative influence on participants’ chunking behavior. The study protocol was approved by the ethics board of the University of Western Ontario.

**Figure 1.**
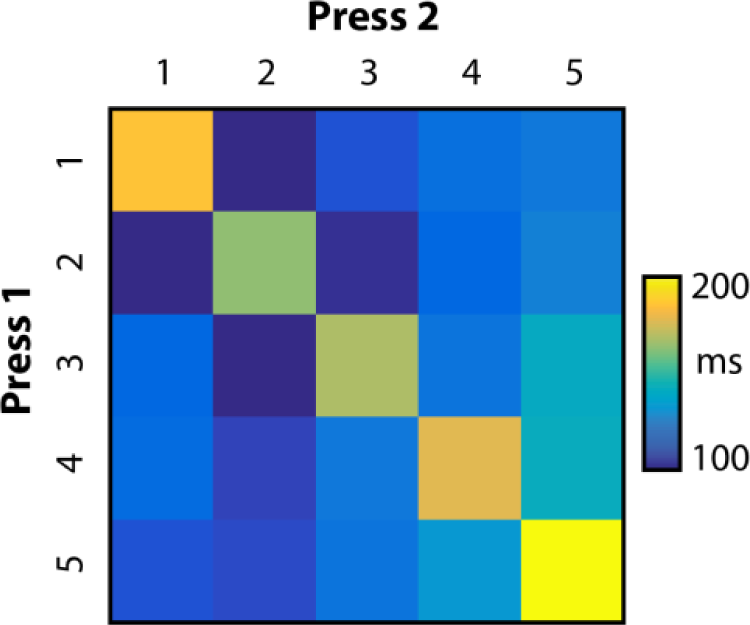
Two-finger transition execution speed. Biomechanical data from an independent dataset in which participants performed all possible combinations of 2 and 3-digit transitions. Matrix indicates the median inter-press interval (IPI) to produce the transition between pairs of keypresses. Indicated values are means over n=7 participants.

### Apparatus

A custom-built five-finger keyboard was used. The keys were not depressible but were equipped with a force transducer (FSG-15N1A, Sensing and Control, Honeywell) underneath each key which measured participants’ isometric force production with a repeatability of <0.02 N and a dynamic range of 16 N (Wiestler and Diedrichsen, 2013; Wiestler et al., 2014; Yokoi et al., 2017). The measured force at each key was digitally sampled at 200 Hz.

### Discrete sequence production task

We used a discrete sequence production task (DSP) in which participants executed sequences of 2, 3, or 11 keypresses as fast as possible while keeping their error rate under 15%. Each trial started with the visual presentation of the sequence to be executed and was completed once the participants pressed the amount of presented numbers. Each block consisted of 28 trials. A trial was deemed erroneous if participants pressed a wrong key anywhere within the sequence. No pause between presses was required and thus some co-articulation between fingers emerged with faster execution. A keypress was registered when the measured force first exceeded 3N. A key release was marked when the force measured at the same key first fell below 1.5N. To prevent participants from pressing more than 2 keys at once, we implemented a constraint such that in order for a key to be registered as depressed the key previously registered as depressed had to be released.

The magnitude of the force applied to each key was represented by 5 lines on an LCD monitor, with the height of the line representing the force at the corresponding key. A white asterisk (memory-guided conditions) or a digit (cued condition) for each finger press was presented above the lines. Immediately after the keypress threshold was reached, participants received visual and auditory feedback. If the correct key was pressed, the color of the cue changed from white to green and a sound was presented. If the incorrect key was pressed, the cue turned red and a lower-pitch sound was presented.

After each trial participants received points based on their accuracy and movement time (MT; the time between the first keypress and last key release). Correct sequences performed faster than the MT threshold (see below) were rewarded with 1 point. MTs that were 20% faster than the threshold were rewarded with 3 points. Incorrect presses or MTs exceeding the threshold resulted in 0 points. At the end of each block, participants received feedback on their error rate, median MT, points obtained during the block, and total points obtained during the session. In order to motivate participants to continue to improve their performance, we adjusted the MT threshold by lowering it by 500 ms after each block in which the participants performed with an error rate of 15% or lower and had a median MT faster than the current threshold. This manipulation resulted in an approximately stable overall error rate of 14.6%, SD: 2.6%. On 27% of trials, participants received 1 point, on 34% of trials 3 points.

### Biomechanical baseline study

To design the chunks and sequences for the main experiment, we conducted a separate study to determine the influence of biomechanical constraints on finger transition speed. 7 participants (5 females, ages: 21-27) participated in this 3-day study. Participants executed all possible two-finger transitions (e.g. 25) and three-finger transitions (e.g. 125), each 8 times per day. Each sequence was presented twice in a row. Each day, participants completed 8 blocks with 150 trials each. The setup and motivational structure were the same as reported above. We found that on our device, transitions between two adjacent fingers (e.g. 12) could be performed faster than two repeated presses of the same finger (e.g. 55; *t*_(6)_ = 13.965, *p* = 8.404e-06; Fig. 1). Given that the 2-3 press sequences hardly taxed the cognitive system, these results can be taken as a characterization of the biomechanical constraints of our specific task. To press the same finger twice, the force applied to the key had to first exceed the press threshold, then go below the release threshold and then cross the press threshold again. This rapid alternation of forces takes time to produce. In contrast, for two adjacent fingers, the second finger press can be initiated (have already reached the press threshold) before the previous finger is released, making it easier to rapidly produce this force pattern. Even though participants improved their overall speed from 157 ms on the first day to 114 ms on the third day, the 5×5 pattern of relative inter-press interval (IPI) was stable both across participants (average correlation *r* = 0.689) and days (*r* = 0.894).

### Experimental design

To experimentally impose a particular way of chunking, we instructed participants in the experimental group to memorize and perform a set of 2-3 keypress chunks (Fig. 2a). These chunks were later combined to form the training sequences (Fig. 2b). Our goal was to impose beneficial or detrimental motor habits on participants’ performance. For this, we used the finding from the biomechanical baseline study that finger repetitions are performed slower than presses of adjacent fingers. We designed sequences such that they would include both fast transitions (runs e.g. 123) and slow finger repetitions (e.g. 113). Depending on which chunk structure was instructed, these transitions would either fall on a chunk boundary or lie within a chunk. In the “aligned” chunk structure we aligned the boundaries such that they fell on difficult finger transitions, which were executed slowly for biomechanical reasons. The time required to perform these difficult finger transitions can therefore simultaneously be used to recall the next chunk, which should benefit overall performance. Using this chunk structure, the 3-digit “runs” (i.e. 123) which are performed quickly were kept intact (not broken up by a chunk transition). We predicted that learning the sequence using this chunk structure would be beneficial to performance speed (Fig. 2c). In the misaligned chunk structure, we placed chunk boundaries in a way that divided up biomechanically easy finger transitions such as runs, thereby breaking up parts of the sequence that could otherwise be performed very quickly. We hypothesized that this would hinder overall performance (Fig. 2c). All participants practiced the same 7 sequences (Fig. 2b). Half of the participants were instructed with the aligned chunk structure for the first 3 sequences, and the misaligned chunk structure for the next 3 sequences (Fig. 2d). For the other half of the participants, the assignment of sequences to aligned and misaligned was reversed. The last sequence served as a control sequence and was chunked, such that either instruction should lead to similarly beneficial performance. The counterbalanced design (Fig. 2d) allowed us to draw strong inferences about whether participants’ performance was dictated by biomechanical demands (which were identical across participants) or whether it was affected by the chunk structure imposed during the induction phase (which was different between the two chunk sets).

**Figure 2.**
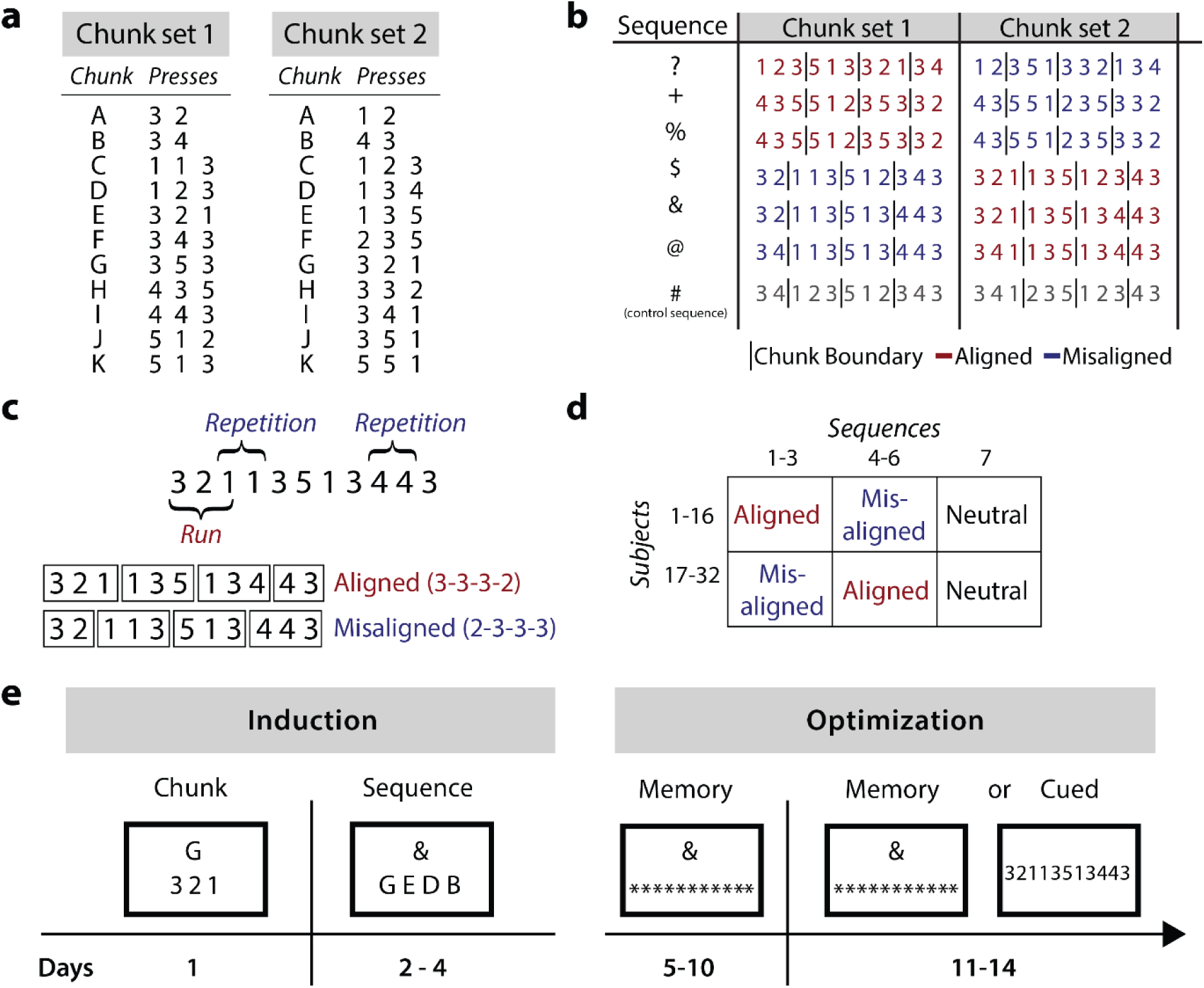
Experimental design. **(a)** Each participant learned 11 chunks associated with the chunk cues (A-K) from one of the chunk sets. **(b)** The seven 11-digit sequences that participants trained on. The vertical lines (not shown to the participants) indicate the chunk boundaries induced in training through the chunk set. Sequences were trained with an aligned (red) or misaligned (blue) chunk structure. **(c)** Example sequence containing a 3-digit run and two-digit repetitions. In the aligned structure, the chunk boundaries fell between repetitions, in the misaligned structure the chunk boundary broke up the run. **(d)** We counterbalanced across participants which sequences were practiced with which chunk structures. **(e)** Experimental timeline depicting the training at each stage. In the induction phase participants memorized chunks and sequences. In the optimization phase participants trained to perform these sequences as fast as possible from memory. In the last week of training, half of the participants were directly cued with the sequence, while the others performed the sequences from memory.

Every participant completed 15 training sessions in total (Figure 2e): one session per day across a 3-week period. Each session lasted approximately 1 hour, excluding the two initial sessions and the last session which each took 2 hours. Participants completed at least 10 blocks of 28 trials per training day. Each block comprised 4 repetitions of each of the 7 sequences.

### Days 1-4: Chunk induction & initial sequence learning

#### Experimental group

On Day 1 the participants were pre-trained on one of the chunk sets (Fig. 2a). Each chunk was associated with a letter of the alphabet (A-K). Participants were explicitly told to learn this association. Each letter A-K was presented twice in succession. In half of the blocks, on the first trial of each pair, the numbers corresponding to the finger presses accompanied the letter on the screen, while on the second trial participants had to recall the presses solely based on the letter (numbers were replaced with stars). This trial order was reversed on every second block. To ensure that participants had memorized the chunks we added speeded recall blocks at the end of days 1 and 2. At the end of the first day, participants could reliably produce the chunks from memory with an average accuracy of 92.7%.

On day 2 participants trained on the seven 11-press sequences. Each sequence was associated with a symbol (e.g. $; suppl. Fig. 2b). Each symbol was presented twice in succession and participants had to perform the sequences from memory using the symbol cue on one trial or with the help of the chunk letters on the next trial. We tested participants’ sequence knowledge with a self-paced recall block at the end of days 2-4 (The first two participants did not perform the recall blocks). At the end of day 4, participants were able to recall all sequences from memory using the sequence cues with an accuracy of 93.1%.

#### Control group

The control group did not receive any chunk training but instead trained directly on the seven 11-press sequences. On day 1 they were presented with the 11 digits corresponding to the 11-press sequences. We matched the amount of training across groups by ensuring that all participants were required to produce the same overall number of finger presses. On day 1, the control participants were not aware that they would have to memorize the sequences later on. On days 2-4 they were instructed to memorize the sequences using the same symbolic sequence cues as the experimental groups and their memory was tested using recall blocks at the end of each day (Day 4: 90.2% accuracy). The rest of the experimental design was identical for all groups.

#### Days 5-10: Optimization - Memory Recall

On days 5-10 participants practiced exclusively on the eleven-press sequences using the symbolic cues. Chunks were no longer cued. Each sequence cue was presented twice in succession and participants had to recall the sequence from memory on both trials.

#### Days 11-14: Optimization - Memory recall or cued presentation

On the last four days of training half of the experimental participants performed the sequences from memory (as on days 5-10), while for the other half and for the control participants we removed the symbolic sequence cue and instead visually presented participants with the complete set of 11 digits that corresponded to the sequences (Fig. 2e). Participants completed an additional generalization test on day 15. The results of this test are not reported in this article.

### Statistical Analysis

We recorded and analyzed the force measured at each key. For each trial, we calculated movement time (MT, time between the first press and last release) and inter-press-intervals (IPIs; time between force peaks of two consecutive presses). All analyses were performed using custom-written code in MATLAB (The MathWorks). We excluded from our analyses trials that contained one or more incorrect presses, as well as trials with an MT or a press with an IPI three standard deviations above the mean calculated across all days and participants. The data were analyzed using mixed-effects analysis of variance (mixed ANOVA), Pearson’s correlation and paired and one-sample t-tests. All t-tests were two-sided. A probability threshold of p<0.05 for the rejection of the null hypothesis was used for all statistical tests. For the regression analyses as well as for calculating the MT difference between the sequences with misaligned and aligned instruction we subtracted the mean performance for each participant and day (across sequences) to normalize and remove the large part of the variance due to interindividual performance differences.

#### Probabilistic model for estimating chunk structure

To estimate participants’ chunking behavior from IPIs, we used an extended version of a Bayesian model of chunking behavior, developed by Acuna et al. (2014). The algorithm uses a Hidden Markov Model to estimate the posterior probability that a specific chunk structure is present on a given trial. Here we used only the IPIs on correct trials, but not the error probability as in the original publication, as the probability of errors did not relate systematically to the imposed chunk structure early in learning.

As we had 10 digit transitions, each of which could either coincide with a chunk boundary or not, we had to consider 2^10^-1= 1023 possible chunk structures. Between trials, the hidden Markov process could either preserve the same chunk structure with probability *p* or switch to any other chunk structure with probability (1-*p*)/1022. The IPIs were modeled as a Gaussian random variable, with a different mean and variance depending on whether the keypress transition was within or between chunks.

In contrast to Acuna et al., in which learning effects were removed in a preprocessing step using a single exponential, we modeled learning within our model using two separate exponential terms for the IPI mean. This captured the faster reduction in the between-compared to the within-chunk intervals (Fig. 3a). The inclusion of separate learning curves for within- and between-chunk IPIs allowed us to estimate participants’ chunk structure independently of changes in the overall performance speed (Fig. 5a). This is an important advance over previous methods that used a constant cutoff value to distinguish between within- and between-chunk intervals. For these methods, faster performance would automatically decrease the number of chunk boundaries detected. To confirm that our algorithm did not show this bias, we simulated artificial data using parameter estimates for individual participants. We simulated sequences that switched between 4 different chunk structures, each of which contained 4 chunks. Even though IPIs decreased by about 300 ms with learning, the estimated average number of chunks remained stable across the entire simulated experiment (average distance to single chunk: 3.35 ∼ 4 chunks and 3 boundaries).

**Figure 3.**
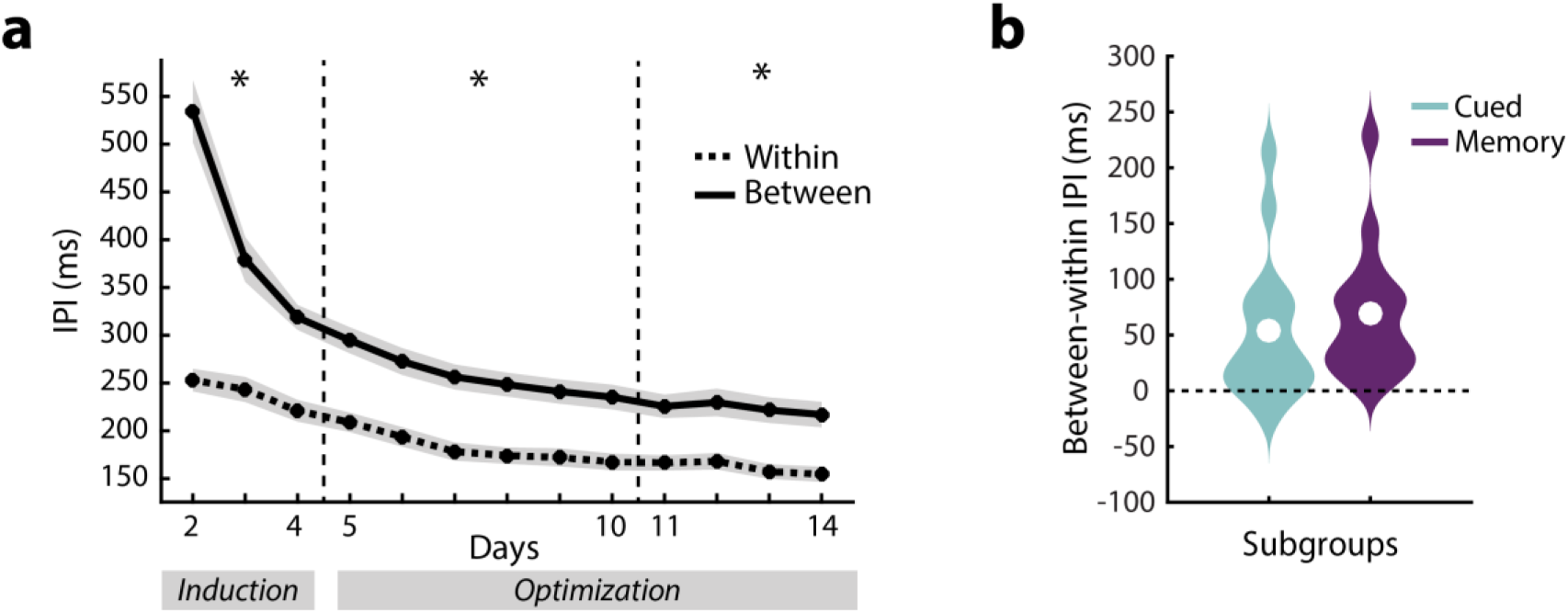
Within- vs. between-chunk inter-press intervals (IPIs). **(a)** Time course of IPIs that were within an instructed chunk (dashed line), or on the boundary between chunks (solid line). Asterisks indicate significant differences between average within- and between-chunk IPIs in the corresponding week (separated by dashed lines). Shaded area denotes between-subject standard error. **(b)** Difference of between- and within-chunk IPIs in the last week of training, split by whether participants had to recall the sequences from memory or were cued with the sequence numbers. Violin plots indicate distribution of individual participants, white circles indicate means.

We used an Expectation-Maximization algorithm to simultaneously estimate the posterior probability of each chunk structure for each trial, as well as the 9 parameters of the model: 3 parameters each for the exponential curve for the within- and between-chunk IPIs, 1 variance parameter for each, and the transition probability *p (for implementation details, see* https://github.com/jdiedrichsen/chunk_inference.

As a preprocessing step, we regressed the IPIs for each subject against the average biomechanical profile, which was estimated as the average IPI profile for all possible 2-digit presses from our biomechanical baseline experiment (Fig. 1). The fitted values were removed from the IPIs. Removing temporal regularities that could be modeled with biomechanics alone should result in chunking estimates that more closely reflect cognitive and learning influences. Qualitatively comparable results were also obtained using the raw IPIs, without biomechanical factors removed.

#### Expected distance

We quantified how much participants changed their chunking behavior over time by calculating the expected distance between their estimated chunk structure and a reference chunk structure. We defined the distance between two chunk structures, *d(i,j)*, as how many of the 10 keypress transitions would have to change from a chunk boundary to a non-boundary (and vice versa) to transform one structure into the other (for an example, see Fig. 5b). A distance of 0 would indicate no change. The average distance between two randomly chosen chunk structures is 5. Because chunk structures produced by participants on each trial were estimates, we calculated the expected distance. For this, we first calculated a 1023 × 1023 matrix containing the distances between any chunk structure *i*, and chunk structure *j*. From the posterior probability distribution, we could then derive how likely each of these chunk structure changes was, *p(i,j)*. The expected value of the distance was then calculated as

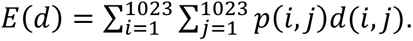

### Data availability

The datasets generated and analyzed during the current study will be made available upon publication.

## Results

Over 15 days we trained 32 participants to produce sequences of 11 isometric keypresses from memory on a keyboard-like device. Participants were rewarded with points for executing sequences as fast as possible while keeping the proportion of incorrect keypresses in each block of trials below 15%. We maintained the participants’ motivation by gradually decreasing the movement time (MT) threshold at which they received points.

We manipulated how participants memorized the sequences by splitting the sequences into several chunks, each composed of 2-3 keypresses. The aim was to test whether the different ways of chunking (hereafter “chunk structures”) imposed through the chunk training in the induction phase (Methods, Fig. 2b) would affect performance optimization in the subsequent two weeks of training. Each sequence could be chunked in an aligned or misaligned fashion, predicted to lead to beneficial or detrimental performance respectively (Methods, Fig. 2c). All participants practiced the same 7 sequences but differed in the chunking instructions they received for each sequence.

### Chunk induction induces a stable motor pattern

To assess whether the imposed chunk structures influenced participants’ motor behavior, we examined inter-press time intervals (IPIs). An increased IPI is commonly taken as a sign of a chunk boundary, as the cognitive processes (memory recall, action selection) involved in switching from one chunk to another require additional time (Verwey, 1999; Verwey et al., 2010). Hence, we would expect our participants to exhibit shorter IPIs between keypresses that belonged to a chunk imposed during day 1 (within-chunk IPIs) and larger IPIs for the boundaries between chunks (between-chunk IPIs). For this analysis, we pooled the data from all sequences irrespective of instruction (misaligned vs. aligned). We indeed found significantly longer between-chunk IPIs compared to within-chunk IPIs in the first few days of training (Fig. 3a: days 2-4: *t*_(31)_ = 7.728, *p* = 5.098e-09), suggesting that our manipulation was successful in inducing a temporally specific pattern of keypresses.

In the optimization phase, we ceased to cue sequences using the alphabetic letters associated with the chunks. Instead, participants were asked to recall the entire 11-keypress sequences from memory in response to symbolic sequence cues (e.g. “$”). Across days 5-10, the within and between-chunk IPIs were still significantly different from each other; *t*_(31)_ = 7.165, *p* = 2.351e-08 (Fig. 3a). This difference cannot be attributed to biomechanical difficulty of the finger transitions, as the within-chunk IPIs for one half of the participants were the between-chunk IPIs for the other half and vice versa (Fig. 2b). IPIs that were within-chunk for all participants (e.g. the first and last IPI of a sequence) were excluded from this analysis. In summary, even though after day 4 we cued the sequences only with symbols, participants persisted in performing the sequences consistent with the chunk structures that we experimentally imposed early in training.

In the last four days of training, we tested whether the persistence of the imposed chunk structure reflected a motor habit or whether it reflected memory recall. Half of the participants continued to perform the sequences from memory, while the other half were cued using the numbers that indicated the necessary keypresses (Fig. 2e), therefore removing any memory recall demands. Both the memory (*t*_(15)_ = 4.865, *p* = 2.059e-04, Fig. 3b) and the cued subgroup (*t*_(15)_ = 3.403, *p* = 0.004) showed a significant difference between the within- and between-chunk IPIs and there was no reliable difference between the two subgroups in this effect (*t*_(30)_ = -0.749, *p* = 0.460). Thus, removing the requirement for memory recall did not abolish chunking. Because none of the subsequent analyses showed any significant difference between the two subgroups, we will report their combined results for the remainder of the article. Overall, these results suggest the explicit chunk training early in learning established a stable performance pattern that outlasted 10 days of subsequent practice.

### Misaligned chunk structure impairs performance

To show that the initial instruction led to the emergence of a motor habit, we needed to not only show that this initial instruction induced a stable temporal pattern of IPIs, but also that this pattern was maintained even when it leads to slower execution speeds than other patterns. We therefore designed chunk structures that were predicted to be either beneficial or detrimental to performance (aligned vs. misaligned respectively) based on their biomechanical constraints (see Methods). Each participant learned 3 of the 7 sequences with a misaligned chunk structure and 3 sequences with an aligned chunk structure, with the assignment counterbalanced across participants (Fig. 2d). This counterbalanced design allowed us to compare execution speed between aligned and misaligned sequences for each participant.

To test our prediction that training with the misaligned chunk structure would lead to poorer performance, we measured participants’ movement time (MT) by estimating the time between the first finger press and the last finger release. In the induction phase, sequences instructed with the misaligned chunk structure were performed slower than the sequences instructed with the aligned chunk structure (Fig. 4a) one-sample t-test: *t*_(31)_ = 2.693, *p* = 0.006). Hence, we were not only able to manipulate how participants performed a sequence, but also how well they could perform it.

**Figure 4.**
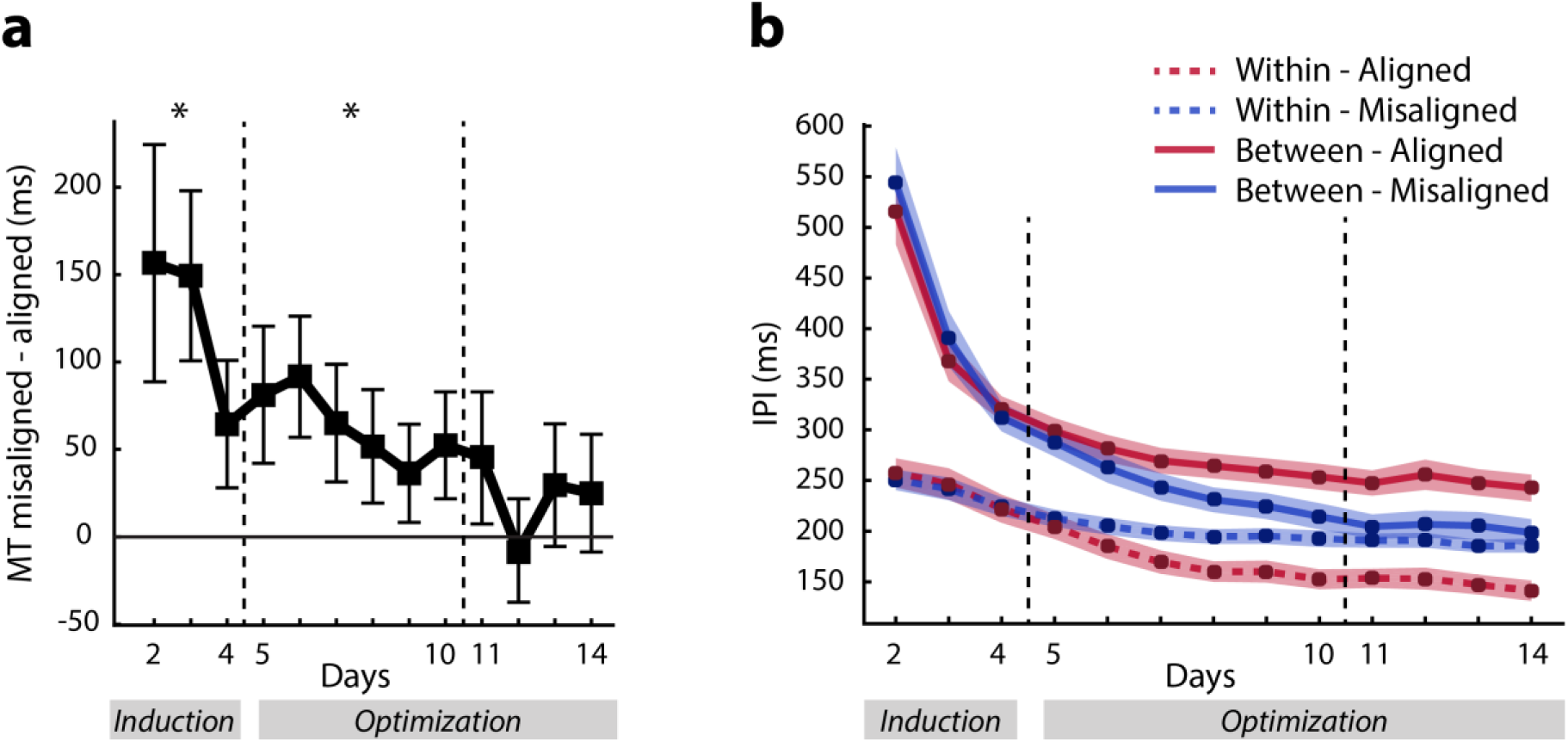
Change in chunk structure and performance for aligned and misaligned instructed sequences. **(a)** Differences in movement time (MT) between sequences instructed with an aligned or misaligned chunk structure. Asterisk indicates a significant difference from 0 (no difference). **(b)** Within-or between-chunk IPIs across training days, separated by whether they were in the aligned or misaligned instructed sequences. Error bars denote between-subject standard error.

**Figure 5.**
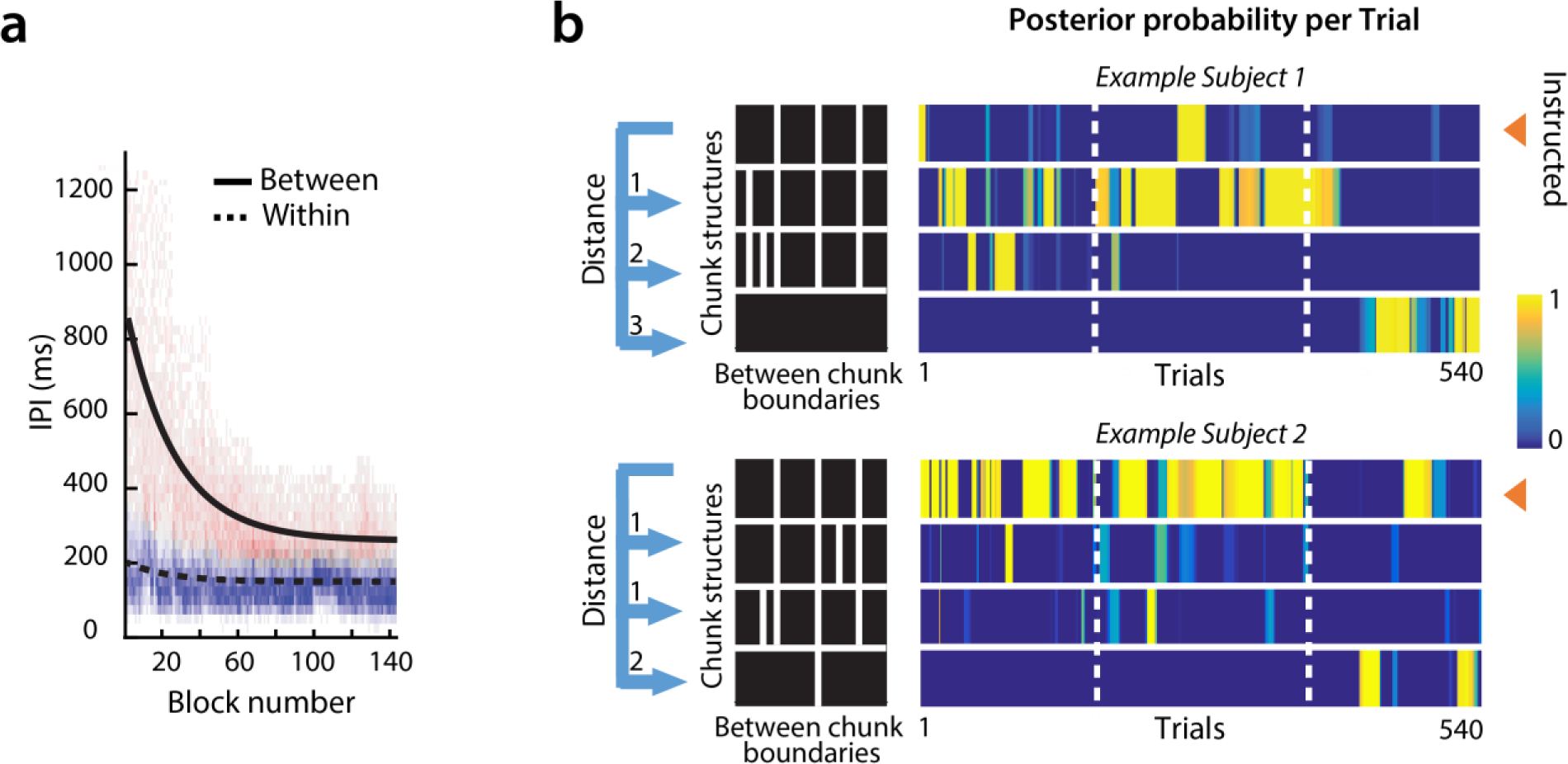
Probabilistic chunking model fitted to example participant data. **(a)** The change of within- and between-chunk IPIs were modeled using two separate exponential functions across training. The density plot shows individual IPIs, with the color indicating the probability of a between-(pink) or within-chunk interval (blue). **(b)** Posterior probability for two example participants (for one sequence per participant) over the course of the experiment. Only the 4 most likely chunk structures out of the 1023 possible structures are shown. The color scale indicates the posterior probability of a given chunk structure for each trial - with yellow indicating higher probabilities. The dashed vertical lines indicate the boundaries between training phases (Days 2-4; 5-10 & 11-14). The black box (left) indicates the chunk boundaries as white lines within the 11-press sequence (max. 10 boundaries) for the chosen chunk structures. The first row indicates the instructed chunk structure (arrow). The other three rows illustrate other chunk structures that were highly probable at some point during the experiment. The distance measure expresses how many chunks need to be added or removed to transform one structure (in this case the instructed chunk structure) into the other.

Examining what factors influenced the difference in speed we observed, we found that on average a within-chunk finger run led to an advantage of 28.6 ms and a within-chunk repetition cost 16 ms. An additional factor that influenced participants’ speed was whether the 2-digit chunk was placed in the beginning (misaligned) or the end of the sequence (aligned), which led to an advantage of 24.7 ms. The difference in MT found in the first week was maintained in the second week of training (days 5-10: *t*_(31)_ = 2.313, *p* = 0.014). Importantly, this shows that the stable pattern of IPIs indeed constitutes a motor habit. This speed difference was no longer statistically reliable in the last four days of training (days 11-14: *t*_(31)_ = 0.764, *p* = 0.225). This suggests that participants were able to overcome the “bad” habit of a misaligned chunk structure to some degree.

### Misaligned chunk structure is changed more rapidly

To investigate how participants overcame the detrimental influence of the misaligned chunk structure, we first separated the IPI analysis (Fig. 3a) by whether the intervals came from sequences that were instructed using an aligned or misaligned structure. While the difference between within- and between-chunk IPIs for sequences constructed using aligned chunk structures was stable over the entire training period, the difference was absent for misaligned chunk structures in the last four days of training (Fig. 4b). The three-way day x within/between x aligned/misaligned interaction was significant (*F*_(12,372)_ = 19.790, *p* = 1e-16). Thus, in the last four days of training participants diverged from the misaligned chunk structure while maintaining the aligned chunk structure.

A disadvantage of this analysis, however, is that we cannot discern how participants restructured their chunking and whether they completely abandoned the misaligned chunk structure. For a clearer understanding of how participants changed their chunk structure, we used a model-based approach to analyze our IPI data.

### Bayesian model of chunk behavior

We used a Bayesian model to estimate the probability of each possible chunk structure given the observed series of IPIs on a trial-by-trial basis (Acuna et al., 2014). The state variable in this Hidden Markov Model represents which of the 1023 possible chunk structures is present on each trial. Using an expectation-maximization (EM) algorithm (Dempster et al., 1977; Welch, 2003), we simultaneously estimated the 9 free parameters of the model (for details see Methods), and the posterior probability for each possible chunk structure on each trial. We accounted for the effects of biomechanical difficulty by regressing out the patterns of IPIs across finger transitions predicted from our biomechanical dataset (Fig. 1) before modeling. Importantly, our model could capture separate learning-related changes to the within- and between-chunk intervals (Fig. 5a). Our method, therefore, allowed us to estimate participants’ chunk structure independently of the overall speed of performance.

Figure 5b shows two examples of individual participants and sequences. In the first panel, the participant chunked the sequence according to the initial instructions at first, then inserted 1 or 2 additional chunk boundaries, and at the end of training performed the sequence as a single chunk. In comparison, the other participant maintained the instructed chunk structure for most of the training period.

To characterize changes in chunk structure over training we defined a metric that quantified the difference between any two chunking structures. The metric is based on counting the number of chunk boundaries that differ, in other words, the number of chunks that would need to be split or merged to transform one chunk structure into the other (Fig. 5b - distance). We then used this measure to calculate, on each trial, the distance between the chunk structure estimated for the participant and three reference structures of interest: (1) the aligned-, (2) misaligned, and (3) a structure that consisted of a single chunk. These distances defined a coordinate system that enabled us to visualize changes in chunk structure over training. We then projected participants’ estimated chunk structures into this space (Fig. 6a). On the horizontal axis is the expected distance of participants’ chunk structure to the single-chunk structure. Given our definition of distance, this measure simply counts the number of chunk boundaries. The vertical axis indicates how close the estimated chunk structure is to the aligned and misaligned chunk structure, respectively.

**Figure 6.**
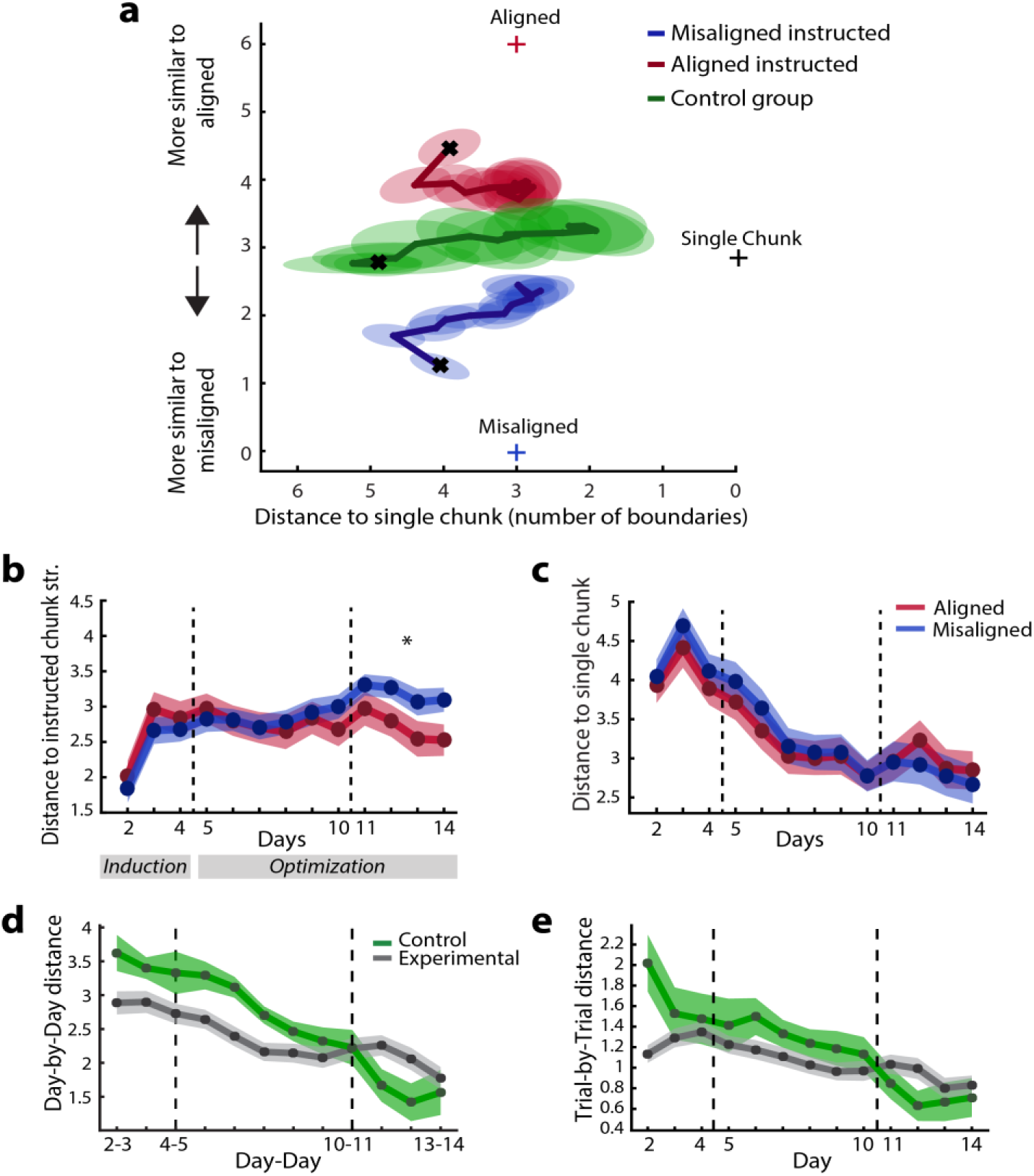
Changes in chunk structure with learning. **(a)** The average chunk structure over 13 days of practice for aligned (red) and misaligned (blue) instructed sequences for the experimental participants. The results of the control group are shown in green. The horizontal axis represents the distance to the single-chunk structure, i.e. the number of chunk boundaries. The vertical axis shows the distance to the aligned or misaligned chunk structure. The crosses indicate the positions of the three reference structures (aligned, misaligned and single). Ellipses denote the between-subject standard error. **(b)** Average distance of participants’ chunk structure to the instructed chunk structure. **(c)** Distance to the single chunk structure across days. **(d)** Day-by-day changes in chunk structure. **(e)** Trial-by-trial changes in chunk structures within each day. Error bars indicate between-subject standard error.

### Participants abandoned the misaligned faster than aligned chunk structure

Using this modeling approach, we probed how much participants diverged from the initial instructions and whether they diverged from the misaligned chunk structure to a greater degree as already suggested by our IPI analysis. Participants slowly changed their chunk structure for both aligned and misaligned instructed sequences with training. The average distance to the instructed chunk structure increased systematically over time (repeated measures ANOVA, effect of day, *F*_(12,372)_ = 7.055, *p* < 1e-16, Fig. 6b).

Consistent with our IPI analysis (Fig. 4b), we observed that participants abandoned the instructed misaligned chunk structure to a greater degree than the aligned chunk structure (Day x Instruction interaction: *F*_(12,372)_ = 5.610, *p* < 1e-16). In the last four days of training, sequences with the misaligned chunk structure were more dissimilar to the instructed chunk structure than sequences with an aligned chunk structure: *t*_(31)_ = 2.294, *p* = 0.029 (Fig. 6b). Additionally, we found a significant Day x Instruction interaction (*F*_(12,372)_ = 2.215, *p* = 0.011) for the distance to a single chunk (Fig. 6c), suggesting a stronger tendency towards performing a sequence as a single chunk when trained on the misaligned chunk structure. Together these results indicate that participants changed their chunking behavior more readily for sequences that were trained using the misaligned chunk structure than when trained using the aligned chunk structure.

Despite the divergence from the misaligned chunk structure with training, our analysis also revealed that participants did not overcome the influence of the instruction completely. In the third week, sequences trained with a misaligned chunk structure were still performed using a chunk structure that was closer to the misaligned structure than to the aligned structure (*t*_(31)_ = 6.962, *p* < 1e-16). This shows that training with a misaligned chunk structure had a lasting influence on participants’ motor behavior.

### Movement towards a single chunk structure

Previous literature has suggested that with training, participants group smaller chunks together to form new larger chunks (Verwey, 1996; Sakai et al., 2003; Kuriyama et al., 2004; Verstynen et al., 2012; Wymbs et al., 2012; Song and Cohen, 2014; Ramkumar et al., 2016), a process that may help to improve performance (Verwey, 1999, 2001; Verwey et al., 2010; Abrahamse et al., 2013; Verwey and Wright, 2014; Ramkumar et al., 2016). However, in nearly all previous studies the estimated number of chunks is biased by the overall movement speed. As verified by simulations (see Methods), our probabilistic model was able to disambiguate the two factors. We estimated the number of chunk boundaries for each participant averaged across sequences (the neutral sequence was excluded). On the 2^nd^ day, participants separated sequences into more chunks than the 4 chunks we instructed (Fig. 6c, *t*_(31)_ = 4.224, *p* = 0.0002). This tendency continued on day 3, on which participants tended to subdivide the sequences into even smaller chunks (day 2 vs. 3: *t*_(31)_ = 2.023, *p* = 0.052). After day three the number of chunk boundaries decreased as shown by a significant effect of day in a repeated measures ANOVA (*F*_(11,341)_ = 11.710, *p* < 1e-16). However, even in the last phase of training, participants performed the sequences with an average of 2.9 chunk boundaries (we instructed 3 chunk boundaries). Thus, while there was a clear tendency towards merging chunks after an initial increase, participants did not perform the sequence as a single chunk, even after 3 weeks of practice.

### Chunk structure crystallizes with training

Would longer training allow participants to completely overcome the influence of the instruction and to perform all sequences as a single chunk? Although experiments with longer training are necessary to provide a definitive answer, our data indicate that this process, if occurring, may take a very long time. The amount of change in the chunk structure for each sequence reduced dramatically in the last week of training, suggesting that a stable motor habit formed. This phenomenon is akin to the development of an invariant temporal and spectral structure in bird-song, a process that has been termed “crystallization” (Brainard and Doupe, 2002). As a measure of crystallization, we calculated the distance between the chunk structures from one day to the next (Fig. 6d) and within each day from one trial to the next (Fig. 6e). The analysis was performed separately for each sequence and participant. Overall, both the day-to-day distance (*F*_(11,330)_ = 18.794, *p* < 1e-16) and the trial-by-trial distance decreased significantly across training days (*F*_(12,456)_ = 13.245, *p* < 1e-16). Therefore, participants appeared to settle onto a stable pattern in the last week. Consequently, additional training would likely only lead to slow changes in their chunk pattern.

In summary, our analyses provide a clear representation of how chunking changes with learning. Firstly, participants diverged from the instructions over time with a quicker deviation from the misaligned chunk structure. Secondly, in line with previous research (Verwey, 1996; Sakai et al., 2003; Kuriyama et al., 2004; Verstynen et al., 2012; Wymbs et al., 2012; Song and Cohen, 2014; Ramkumar et al., 2016) participants gradually moved towards performing the sequence as a single chunk by dividing the sequence into fewer chunks. Nevertheless, they did not completely overcome the initial instruction, nor did they perform the sequences as a single chunk at the end of training. Considering that the chunk structure crystallized in the last four days of training, these results demonstrate the formation of a stable motor habit that is still influenced by the initial instruction.

### Spontaneously emerging chunk structures

To investigate how participants might spontaneously chunk the sequences, we tested an additional control group (N=8), who did not receive any explicit chunk training. Participants were presented with the sequences in entirety on the first day and were asked to memorize them without any reference to chunks (see Methods for details). Even though memorization was more difficult, the control group did not differ significantly from the experimental groups in terms of their explicit knowledge on day 4 (*t*_(36)_ = 1.288, *p* = 0.206), or in their overall MT across training (main effect of group: *F*_(1,38)_ = 0.101, *p* = 0.753; interaction between group and day (*F*_(1,38)_ = 1.387, *p* = 0.168).

Similar to the experimental groups, the control group initially subdivided the sequences into small chunks and then slowly combined them into larger chunks. The distance to a single chunk structure decreased significantly over days (*F*_(12,84)_ = 17.977, *p* < 1e-16, Fig. 6a), and reached a level that was not statistically different from the experimental participants on the last day of training (*t*_(38)_ = -0.940, *p* = 0.353). Interestingly, on the first day, the control group performed the sequences closer to the misaligned chunk structure than to the aligned chunk structure (*t*_(7)_ = -2.799, *p* = 0.027). With training, participants then moved closer to the aligned chunk structure, as indicated by a significant change in the difference between the distance to the aligned and misaligned chunk structure across days (*F(*12,84) = 5.303, *p* < 1e-16). The control group also showed clear crystallization over time (see Figure 6d&e). Compared to the experimental groups, control participants showed a higher day-to-day and trial-by-trial change in the beginning of training, which then reduced more quickly (Group x Day interaction; day-to-day: *F*_(11,330)_ = 3.780, *p* = 4.003e-05; trial-by-trial: *F*_(12,456)_ = 4.254, *p* = 2.167e-06). In summary, the control group showed similar behavioral patterns to the experimental participants, indicating that similar processes of habit formation are also at play in the absence of explicit instructions.

### Idiosyncratic chunk structures at the end of training and their importance to performance

Finally, we analyzed how the final chunk structure that participants adopted for each sequence influenced their performance after 3 weeks of training. We visualized this relationship by plotting the chunk structure for each sequence and participant in the 2-dimensional space defined in earlier Fig. 6a, with the corresponding average MT indicated by the size of the symbol (Fig. 7).

**Figure 7.**
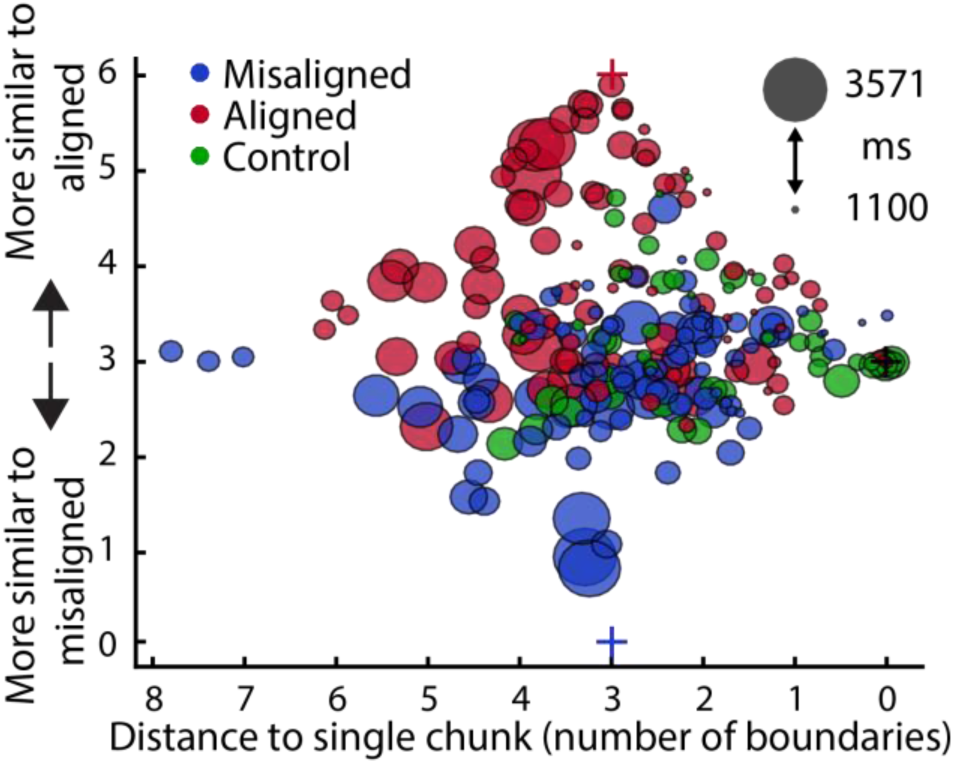
Relationship between chunking and speed (days 11-14). The x-axis indicates the distance to a single chunk and the y-axis the relative distance to the two instructed chunk structures. Each data point indicates the average chunk structure and MT of a single sequence and participant in the last four days of training. The diameter of each circle represents the MT with larger circles indicating slower performance.

The first insight is that participants used quite diverse chunk structures. To show that this is not due to within-subject variability of performance, we compared participants’ within-subject variation in IPI patterns for each sequence across even and odd trials (in the last three days of training) to the between-subject variation in IPI patterns for each sequence. We found that the between-subject variability was much higher than the within-subject variability (t_(31)_ = 36.130, p < 1e-16). This clearly shows that participants developed their own, idiosyncratic way of chunking each sequence, which is not fully dictated by the biomechanical requirements of the sequence. With this result in mind, we asked whether these individual differences relate to differences in final performance.

Figure 7 suggests, that performance was better for sequences that were closer to the aligned chunk structure. To statistically test whether this finding holds true within each individual, we regressed the MT for 6 sequences (last 4 days & excluding the control sequence) for each participant in the last four days of training against the corresponding distance to the aligned chunk structure. On average the individual slopes were significantly greater than 0, both for the experimental (Fig. 8a; *t*_(31)_ = 2.220, *p* = 0.017), and control group (Fig. 8b, *t*_(7)_ = 2.720, *p* = 0.015). Thus, finding a better way of chunking (for the same number of chunk boundaries) improved performance.

**Figure 8.**
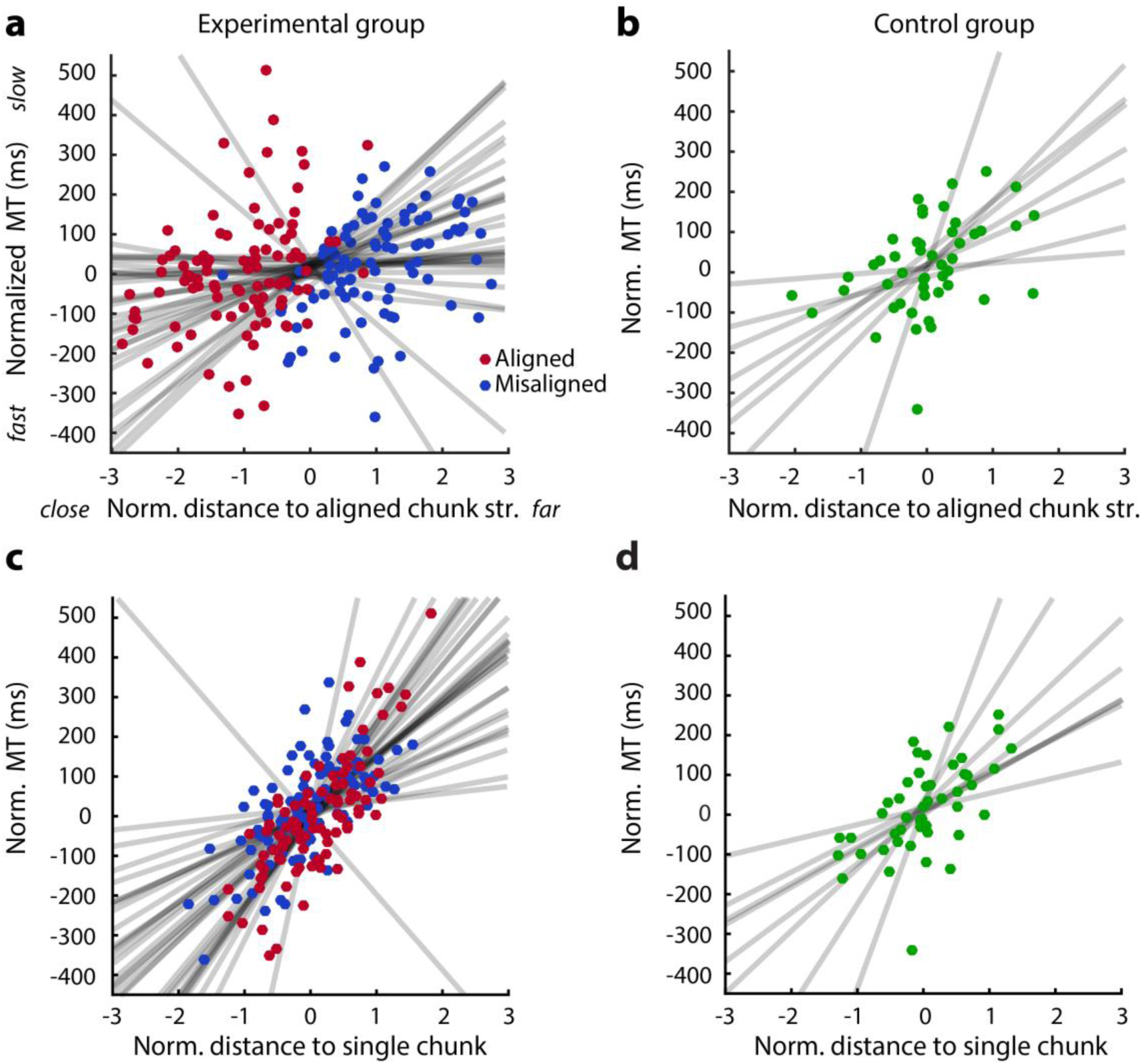
Relationship between the distance to the aligned/single chunk structure and MT. **(a)** Scatterplot between the normalized (per subj.) distance to the aligned chunk structure and normalized MT in the last four days of practice. A separate regression line is fitted to the 6 sequences for each participant. Red dots indicate sequences with aligned instructions, blue dots sequences with misaligned chunking instructions. **(b)** Same as **a** but for the control group. **(c&d)** same as a & b but for the normalized distance to a single chunk.

Secondly, Fig. 7 also suggests, that performing the sequence with a reduced number of chunks is beneficial for performance. We regressed the MT for 6 sequences (last 4 days & excluding the control sequence) against the corresponding distance to the single chunk structure to (Fig. 8c). The majority of the participants showed a positive relationship between the number of chunks and MT: a one-sample t-test indicated that the individual slopes were significantly greater than 0 (*t*_(31)_ = 6.104, *p* = 4.560e-07). This relationship was also found for the control participants (Fig. 8d, *t*_(7)_ = 3.429, *p* = 0.006). Thus, performing the sequences with fewer chunks led to better performance. Note that for both analyses, the chunk structure can be determined independently from the overall performance criterion (MT, see Methods).

Overall, these results suggest that the two optimization processes - joining chunks and aligning the remaining chunk boundaries with biomechanical constraints - positively influenced participants’ ultimate performance. Sequences for which participants could not develop a better way of chunking were performed substantially slower.

## Discussion

In this study, we utilized chunking as a tool to investigate the role of motor habits in skill learning. We influenced the structure of the initial declarative sequence representation by manipulating how participants memorized them (Park, Wilde, & Shea, 2004). By experimentally imposing two different chunk structures on the same physical sequence, we could make causal inferences about the effects of cognitive chunking on motor skill development. This is an important advance over previous observational studies (Wright et al., 2010; Wymbs et al., 2012; Ramkumar et al., 2016), which did not experimentally control how participants chose to chunk sequences.

We report three main results. First, consistent with previous studies (de Kleine & Verwey, 2009; Verwey et al., 2010, 2009; Verwey & Dronkert, 1996), our data demonstrate that a stable chunking pattern can be induced through cognitive manipulations during the initial stages of sequence learning. Importantly, participants did not completely overcome this imposed chunk structure, even after 2 weeks of additional training. Participants’ chunk structure crystallized towards the end of training, making it unlikely that the influence of the initial instruction would disappear completely with longer practice. Finally, the chunking structure remained stable, even when the task changed from a memory-guided to a stimulus-guided task. Thus, the initial instruction led to the formation of specific motor patterns that were still clearly measurable after three weeks of training.

Second, we tested whether this stable pattern of chunking could be considered a motor habit. To do so, we designed two different ways of instructing the sequence, one aligned and the other misaligned with biomechanical influences. This manipulation either facilitated or impeded performance in the first two weeks of practice. We showed that participants did not overcome the misaligned structure completely, even though it was detrimental to their performance. Thus, the stable chunking pattern meets the requirements (as laid out in our definition) for being called a motor habit. Therefore, we believe that studying sequential chunking can provide valuable insights into the neural systems underlying motor habits. Indeed, it has recently been suggested that chunking plays an integral role in the formation and expression of habits (Dezfouli, Lingawi, & Balleine, 2014; Graybiel, 2008) and is neurally represented in the dorsal lateral striatum as action “start and stop signals” (Barnes, Kubota, Hu, Jin, & Graybiel, 2005; Graybiel, 1998; Jin, Tecuapetla, & Costa, 2014; Smith & Graybiel, 2013a, 2014).

Finally, our results also indicate that the motor habit was not completely immutable. Participants were able to modify the misaligned chunk structure and did so more rapidly than the aligned chunk structure. As a consequence, the performance detriment imposed by the misaligned instruction was no longer significant on the group level in the last week of training.

We identified two ways in which participants overcame the limitation induced by the bad habit. After initially breaking up the instructed sequences into 5 chunks on average, participants then joined chunks together, and by doing so, decreasing the amount of additional time spent on chunk boundaries. While previous research has suggested that the size of chunks increases with training, these findings were usually conflated with the overall speed of the action (Wymbs et al., 2012; Song and Cohen, 2014; Solopchuk et al., 2016). Using a Bayesian model to assess chunk structure independent of performance, we demonstrated a positive relationship between chunk concatenation and execution speed, both in the experimental as well as in the control group that developed a chunking strategy without explicit instructions. However, our results also indicate that participants did not merge all sequences into a single chunk after 3 weeks of training, but on average subdivided each sequence into 3-4 chunks. This suggests that the number of motor actions that can be joined in a single chunk may be limited (Verwey et al., 2002; Verwey and Eikelboom, 2003; Langan and Seidler, 2011; Ramkumar et al., 2016).

We found that participants also optimized performance by rearranging chunk boundaries in a biomechanically efficient manner. Consistent with our prediction based on the difficulty of individual digit transitions, placing chunk boundaries at digit transitions that take more time to execute and combining finger presses that are adjacent resulted in faster performance for the full sequence. This optimization process was also observable in the control group that memorized and practiced sequences on their own terms.

Conversely, we observed that sequences that were not chunked in line with these strategies were performed slower. Therefore, if a more beneficial way of chunking was not found, participants still produced sequences using longer movement times, suggesting that other learning mechanisms did not fully make up for a persistent motor habit. Considering that participants’ behavior became highly invariant in the last week of practice, we predict that some motor habit will remain and continue to influence participants’ performance even after prolonged training.

In many motor tasks, there are numerous strategies and processes that can lead to excellent performance (Verwey et al., 2010; Verstynen et al., 2012). Examining Figure 7, one can observe that the shortest MTs were achieved anywhere in the space between the aligned and single chunk structure. Occasionally, good performance was also reached in other locations in chunk space. Our analysis showed that participants adopted quite idiosyncratic chunk structures for each sequence at the end of training. This suggests that there is considerable inter-individual variability in which technique works best for reaching a high level of performance. Part of these differences may reflect biomechanical variation across participants, leading to slightly different optimal solutions. Alternatively, these differences may be learning-related. A number of ways of chunking may work approximately equally well, such that the cost of changing an established habit may outweigh the small benefit that could be gained from changing the structure. A similar observation can be made in sports, where even top-ranked athletes use slightly different techniques to reach similar levels of performance.

The establishment of a novel experimental paradigm to study motor habit formation will allow us to explore ways to encourage learners to abandon or change a current habit. While our attempt at accelerating this process by changing the task from a memory-based to a stimulus-based task was ultimately not successful, there are many other techniques that would be possible. In many disciplines, teachers have developed ways to help students overcome habits. For instance, the Hanon piano exercise helps students play difficult passages of a musical piece by breaking up learned phrases into new chunks to explore different rhythms. Playing a passage slower than intended has also been suggested to break habits (Chang, 2016). Overall, the general advice from the diverse literature on learning piano is to diversify training and to practice with careful awareness to prevent habits from forming (Sadnicka et al., 2017). This suggests that changes in context and the exploration of novel ways of moving can aid performance and the abandonment of habits.

While our experimental design enabled us to manipulate participants’ habits in a laboratory setting, sequence learning only captures a specific aspect of motor skill acquisition. Nevertheless, similar persistence of habits has been observed in other motor learning paradigms (Diedrichsen, White, Newman, & Lally, 2010). In bimanual coordination, for instance, Park, Dijkstra and Sternard (2013) showed that an acquired pattern stayed remarkably stable even over 8 years of not performing the task.

The current study shows that motor habits can be cognitively induced and can remain stable for extended time periods, even though they may prevent further performance gains. Furthermore, the study provides the first insights into potential learning processes that are involved in overcoming a detrimental habit. Our experimental paradigm allows the further study of how we can aid the abandonment of bad habits.

## Acknowledgments

This work is supported by a James S. McDonnell Foundation Scholar award, a Natural Sciences and Engineering Council of Canada (NSERC) Discovery Grant (RGPIN-2016-04890) and the Canada First Research Excellence Fund (BrainsCAN) to J.D., a NSERC Discovery Grant (RGPIN 238338) and a Canadian Institutes of Health Research Grant (PJT-153447) to P.L.G. We thank Aaron L. Wong for helpful comments on earlier versions of this manuscript.

## References

1. Abrahamse EL, Ruitenberg MFL, de Kleine E, Verwey WB (2013) Control of automated behavior: insights from the discrete sequence production task. Front Hum Neurosci 7:1–16.

2. Acuna DE, Wymbs NF, Reynolds CA, Picard N, Turner RS, Strick PL, Grafton ST, Kording KP (2014) Multifaceted aspects of chunking enable robust algorithms. J Neurophysiol 112:1849–1856.

3. Adams CD (1982) Variations in the sensitivity of instrumental responding to reinforcer devaluation. Q J Exp Psychol Sect B 34:77–98.

4. Ashby FG, Ell SW, Waldron EM (2003) Procedural learning in perceptual categorization. Mem Cogn 31:1114–1125.

5. Barnes TD, Kubota Y, Hu D, Jin DZ, Graybiel AM (2005) Activity of striatal neurons reflects dynamic encoding and recoding of procedural memories. Nature 437:1158–1161.

6. Bo J, Seidler RD (2009) Visuospatial working memory capacity predicts the organization of acquired explicit motor sequences. J Neurophysiol 101:3116–3125.

7. Brainard MS, Doupe AJ (2002) What songbirds teach us about learning. Nature 417:351–358.

8. Chang CC (2016) Fundamentals of Piano Practice, 3rd ed. CreateSpace Independent Publishing Platform.

9. Dempster AP, Laird NM, Rubin DB (1977) Maximum likelihood from incomplete data via the EM algorithm. J R Stat Soc Ser B Methodol 39:1–38.

10. Dezfouli A, Balleine BW (2012) Habits, action sequences and reinforcement learning. Eur J Neurosci 35:1036–1051.

11. Dezfouli A, Lingawi NW, Balleine BW (2014) Habits as action sequences: hierarchical action control and changes in outcome value. Philos Trans R Soc B Biol Sci 369:20130482–20130482.

12. Dickinson A (1985) Actions and Habits: The Development of Behavioural Autonomy. Philos Trans R Soc B Biol Sci 308:67–78.

13. Diedrichsen J, Kornysheva K (2015) Motor skill learning between selection and execution. Trends Cogn Sci 19:227–233.

14. Diedrichsen J, White O, Newman D, Lally N (2010) Use-Dependent and Error-Based Learning of Motor Behaviors. J Neurosci 30:5159–5166.

15. Ericsson KA et al. (1993) The role of deliberate practice in the acquisition of expert performance. Psychol Rev 100:363–406.

16. Graybiel AM (1998) The basal ganglia and chunking of action repertoires. Neurobiol Learn Mem 70:119–136.

17. Graybiel AM (2008) Habits, Rituals, and the Evaluative Brain. Annu Rev Neurosci 31:359–387.

18. Graybiel AM, Grafton ST (2015) The Striatum: Where Skills and Habits Meet. Cold Spring Harb Perspect Biol 7:a021691.

19. Haith AM, Krakauer JW (2018) The multiple effects of practice: skill, habit and reduced cognitive load. Curr Opin Behav Sci 20:196–201.

20. Halford GS, Wilson WH, Phillips S (1998) Processing capacity defined by relational complexity: implications for comparative, developmental, and cognitive psychology. Behav Brain Sci 21:803–864.

21. Hardwick RM, Forrence AD, Krakauer JW, Haith AM (2019) Time-dependent competition between goal-directed and habitual response preparation. Nat Hum Behav 3:1252–1262.

22. Hayes JR (2013) The Complete Problem Solver. Taylor & Francis.

23. Hélie S, Waldschmidt JG, Ashby FG (2010) Automaticity in rule-based and information-integration categorization. Attention, Perception, Psychophys 72:1013–1031.

24. Jager W (2003) Breaking ‘ bad habits ’: a dynamical perspective on habit. Hum Decis Mak Environ Percept Underst Assist Hum Decis Mak Real-Life Settings (L Hendrickx, W Jager, L Steg, eds), Lib Amicorum Charles Vlek, Univ Groningen, Groningen, Netherlands:149–160.

25. Jin X, Tecuapetla F, Costa RM (2014) Basal ganglia subcircuits distinctively encode the parsing and concatenation of action sequences. Nat Neurosci 17:423–430.

26. Jog MS, Kubota Y, Connolly CI, Hillegaart V, Graybiel AM (1999) Building Neural Representations of Habits. Science (80-) 286:1745–1749.

27. Kleine E De, Verwey WB (2009) Representations underlying skill in the discrete sequence production task: effect of hand used and hand position. Psychol Res Psychol Forsch 73:685–694.

28. Kuriyama K, Stickgold R, Walker MP (2004) Sleep-dependent learning and motor-skill complexity. Learn Mem 11:705–713.

29. Langan J, Seidler RD (2011) Age differences in spatial working memory contributions to visuomotor adaptation and transfer. Behav Brain Res 225:160–168.

30. Miller GA (1956) The magical number seven, plus or minus two: some limits on our capacity for processing information. Psychol Rev 63:81–97.

31. Moors A, De Houwer J (2006) Automaticity: A Theoretical and Conceptual Analysis. Psychol Bull 132:297–326.

32. Park J-H, Wilde H, Shea CH (2004) Part-Whole Practice of Movement Sequences. J Mot Behav 36:51–61.

33. Park S-W, Dijkstra TMH, Sternad D (2013) Learning to never forget—time scales and specificity of long-term memory of a motor skill. Front Comput Neurosci 7:1–13.

34. Ramkumar P, Acuna DE, Berniker M, Grafton ST, Turner RS, Kording KP (2016) Chunking as the result of an efficiency computation trade-off. Nat Commun 7:12176.

35. Robbins TW, Costa RM (2017) Habits. Curr Biol 27:R1200–R1206.

36. Sadnicka A, Kornysheva K, Rothwell JC, Edwards MJ (2017) A unifying motor control framework for task-specific dystonia. Nat Rev Neurol 14:116–124.

37. Sakai K, Kitaguchi K, Hikosaka O (2003) Chunking during human visuomotor sequence learning. Exp Brain Res 152:229–242.

38. Seger CA, Spiering BJ (2011) A critical review of habit learning and the Basal Ganglia. Front Syst Neurosci 5:1–9.

39. Seidler RD, Bo J, Anguera JA (2012) Neurocognitive contributions to motor skill learning: The role of working memory. J Mot Behav 44:445–453.

40. Smith KS, Graybiel AM (2013a) Using optogenetics to study habits. Brain Res 1511:102–114.

41. Smith KS, Graybiel AM (2013b) A dual operator view of habitual behavior reflecting cortical and striatal dynamics. Neuron 79:361–374.

42. Smith KS, Graybiel AM (2014) Investigating habits: strategies, technologies and models. Front Behav Neurosci 8:1–17.

43. Smith KS, Graybiel AM (2016) Habit formation coincides with shifts in reinforcement representations in the sensorimotor striatum. J Neurophysiol 115:1487–1498.

44. Solopchuk O, Alamia A, Olivier E, Ze A, Zénon A (2016) Chunking improves symbolic sequence processing and relies on working memory gating mechanisms. Learn Mem 23:108–112.

45. Song S, Cohen L (2014) Impact of conscious intent on chunking during motor learning. Learn Mem 21:449–451.

46. Verstynen T, Phillips J, Braun E, Workman B, Schunn C, Schneider W (2012) Dynamic Sensorimotor Planning during Long-Term Sequence Learning: The Role of Variability, Response Chunking and Planning Errors Balasubramaniam R, ed. PLoS One 7:e47336.

47. Verwey WB (1996) Buffer loading and chunking in sequential keypressing. J Exp Psychol Hum Percept Perform 22:544–562.

48. Verwey WB (1999) Evidence for a multistage model of practice in a sequential movement task. J Exp Psychol Hum Percept Perform 25:1693–1708.

49. Verwey WB (2001) Concatenating familiar movement sequences: The versatile cognitive processor. Acta Psychol (Amst) 106:69–95.

50. Verwey WB, Abrahamse EL, de Kleine E (2010) Cognitive processing in new and practiced discrete keying sequences. Front Psychol 1:32.

51. Verwey WB, Abrahamse EL, Jiménez L (2009) Segmentation of short keying sequences does not spontaneously transfer to other sequences. Hum Mov Sci 28:348–361.

52. Verwey WB, Dronkert Y (1996) Practicing a Structured Continuous Key-Pressing Task: Motor Chunking or Rhythm Consolidation? J Mot Behav 28:71–79.

53. Verwey WB, Eikelboom T (2003) Evidence for Lasting Sequence Segmentation in the Discrete Sequence-Production Task. J Mot Behav 35:171–181.

54. Verwey WB, Lammens R, Honk J Van (2002) On the role of the SMA in the discrete sequence production task: a TMS study. Neuropsychologia 40:1268–1276.

55. Verwey WB, Wright DL (2014) Learning a keying sequence you never executed: Evidence for independent associative and motor chunk learning. Acta Psychol (Amst) 151:24–31.

56. Welch LR (2003) Hidden Markov Models and the Baum-Welch Algorithm. IEEE Inf Theory Soc Newsl 53:1,10–13.

57. Wickens JR, Horvitz JC, Costa RM, Killcross S (2007) Dopaminergic Mechanisms in Actions and Habits. J Neurosci 27:8181–8183.

58. Wiestler T, Diedrichsen J (2013) Skill learning strengthens cortical representations of motor sequences. Elife 2:1–20.

59. Wiestler T, Waters-Metenier S, Diedrichsen J (2014) Effector-Independent Motor Sequence Representations Exist in Extrinsic and Intrinsic Reference Frames. J Neurosci 34:5054–5064.

60. Wong AL, Lindquist MA, Haith AM, Krakauer JW (2015) Explicit knowledge enhances motor vigor and performance: motivation versus practice in sequence tasks. J Neurophysiol 114:219–232.

61. Wright DL, Rhee J-H, Vaculin A (2010) Offline Improvement during Motor Sequence Learning Is Not Restricted to Developing Motor Chunks. J Mot Behav 42:317–324.

62. Wymbs NF, Bassett DS, Mucha PJ, Porter MA, Grafton ST (2012) Differential Recruitment of the Sensorimotor Putamen and Frontoparietal Cortex during Motor Chunking in Humans. Neuron 74:936–946.

63. Yokoi A, Bai W, Diedrichsen J, Yokoi XA, Bai W, Diedrichsen XJ (2017) Restricted transfer of learning between unimanual and bimanual finger sequences. J Neurophysiol 117:1043–1051.

